# Analysis of Circulating Exosomes Reveals a Peripheral Signature of Astrocytic Pathology in Schizophrenia

**DOI:** 10.1101/2020.02.18.955013

**Authors:** Mohini Ranganathan, Mohamed Rahman, Suhas Ganesh, Deepak C D’Souza, Patrick D Skosnik, Rajiv Radhakrishnan, Surbhi Pathania, Thalachallour Mohanakumar

## Abstract

**Background:** There is considerable interest in identifying peripheral biomarkers that reflect neuropathological changes in schizophrenia. Extracellular vesicles, including exosomes can cross the blood brain barrier with their contents intact and their cargo, including lipids, proteins and genetic materials can be assayed peripherally. Circulating exosomes have been studied in other neurodegenerative disorders, but there is scarce data in schizophrenia.

**Methods:** We examined neuropathology relevant protein biomarkers in circulating exosomes from patients with schizophrenia and age and gender matched healthy controls using methods consistent with the recommended “Minimal information for studies of extracellular vesicles 2018” (MISEV2018) guidelines. Nanoparticle tracking analysis was used to determine the size and concentration of exosomes. Exosomal membrane marker (CD9) and specific target cargo proteins (glial fibrillary acid protein[GFAP], synaptophysin and α-II-spectrin) immunopositivity was examined using Western blot analyses and band intensity quantified.

**Results:** No group differences were observed between schizophrenia and control samples in plasma exosomal concentration and size or in CD9 or calnexin positivity. Exosomal GFAP concentration was significantly higher and α-II-spectrin expression significantly lower in schizophrenia samples and there were no group differences observed for exosomal synaptophysin concentration.

**Conclusions:** Our results demonstrate for the first time, a differential pattern of exosomal protein expression in schizophrenia compared to matched healthy controls, consistent with the hypothesized astro-glial pathology in this disorder. These results warrant further examination of circulating exosomes as vehicles of novel peripheral biomarkers of disease in schizophrenia and other neuropsychiatric disorders.

## Introduction

Schizophrenia, a devastating illness affecting approximately 1% of the population, has onset of symptoms in adolescence or early adulthood, a heterogenous symptom profile and significant functional impairments (Kahn et al., 2015). Most of what is known about the neuropathology of schizophrenia is from studies of postmortem tissues. The search for peripheral biomarkers that reflect neuropathological changes in schizophrenia is ongoing, but obtaining tissues informative of neuropathology in living patients is an obstacle and results have remained elusive. The study of circulating extracellular vesicles assayed peripherally is one promising approach to identify putative peripheral biomarkers of schizophrenia.

Extracellular Vesicles (EVs) include several distinct types of cell-derived vesicles that are released into the extracellular environment and influence adjacent as well as remote physiological and pathological processes (Colombo, Raposo, & Thery, 2014; Mager, 2020; Saeedi, Israel, Nagy, & Turecki, 2019; Stahl & Raposo, 2019). Exosomes refer to specific EVs that are typically 30-200nm in diameter and arise from late endosomes or multi-vesicular bodies (MVB) (Lee, El Andaloussi, & Wood, 2012). They are secreted into the extracellular space via a distinct secretory pathway and are detectable in biological fluids (Colombo et al., 2014; Stahl & Raposo, 2019).

Exosomes can be secreted from a wide range of cell types into the extracellular space and can be assayed from biological fluids (Thery et al., 2018). In the central nervous system (CNS), exosomes may be secreted by neurons, astrocytes, microglia and oligodendrocytes and appear to be involved in several physiological processes such as neuron-neuron or glial-neuron communication, regulation of myelination, regulation of stress and immune responses and modulation of synaptic plasticity (Mager, 2020; Stahl & Raposo, 2019). Exosomes cross the blood brain barrier, bidirectionally, in health and in illness, with their contents intact (Saeedi et al., 2019), that makes them particularly attractive vehicles of biomarkers for neuropathology.

Analyses of exosomal content provides information about the cell of origin, the inflammatory state of the cellular milieu, and ongoing physiological or pathological processes. The nucleic acid content, in particular, the microRNA content may help elucidate downstream cellular targets and potentially, the functional impact of the exosomes. Post mortem studies have shed light on the role of exosomes in the pathogenesis and disease progression of several neurodegenerative disorders (Levy, 2017) and the ability to identify markers of neuropathology in exosomes obtained from blood or CSF offers great promise in the development of peripheral biomarkers. The study of exosomes in neuropathology has largely been limited to disorders such as Alzheimer’s disease, Parkinson’s disease, Lewy body dementia and traumatic brain injury (TBI) (Goetzl et al., 2015; Goetzl et al., 2019; Goetzl, Kapogiannis, et al., 2016; Goetzl, Mustapic, et al., 2016; Goetzl, Schwartz, Abner, Jicha, & Kapogiannis, 2018; Gruzdev, Yakovlev, Druzhkova, Guekht, & Gulyaeva, 2019; Levy, 2017; Winston, Goetzl, Schwartz, Elahi, & Rissman, 2019). These promising results have provided the impetus to search for similar biomarkers in psychiatric disorders such as schizophrenia.

Thus far, only a handful of published studies have examined exosomes in schizophrenia. Banigan et al (Banigan et al., 2013) analyzed exosomes extracted from frozen post mortem brain tissue of patients with schizophrenia and bipolar disorder and healthy controls and reported greater expression of disease specific exosomal miRNAs in patients, specifically miR-497 in schizophrenia and miR-29c in bipolar disorder. More recently, Amoah et al (Amoah et al., 2019) also examined exosomal microRNA in the orbitofrontal cortex of port mortem brains and reported increased expression of miR-223, that targets glutamate receptors, in schizophrenia and bipolar disorder compared to controls. Further, miR-223 was found to be enriched in astrocytes and sensitive to antipsychotic exposure in this study. Two recent studies have examined plasma exosomes in schizophrenia. Du et al validated several distinct miRNAs in serum exosomes, including 11 that could distinguish between the schizophrenia and the control groups (Du et al., 2019). Finally, Kapogiannis et al reported altered expression of proteins relevant to insulin signaling in plasma derived neuronal exosomes from drug naïve first episode schizophrenia patients, suggesting that altered insulin signaling precedes antipsychotic exposure in schizophrenia (Kapogiannis et al., 2019).

To our knowledge, there are no published studies examining neuropathology relevant proteins in circulating exosomes in schizophrenia.

This cross-sectional study examined circulating exosomes from plasma of patients with schizophrenia and matched healthy controls. Target proteins were identified based on the following criteria 1) cargo proteins previously identified in peripheral blood exosomal assays of disorders with significant neuropathology including dementias and TBI (Karnati et al., 2019; Sun, Dalvi, Abadjian, Tang, & Pulliam, 2017) ; 2) proteins with relative cell specificity to the CNS neurons (axonal and synaptic) and glia (astrocytes); and 3) proteins with vital roles in neurodevelopmental processes, perturbations of which result in Mendelian syndromes with severe neuropsychiatric sequalae. Based on these criteria, synaptophysin, (a neuronal synaptic vesicular membrane protein), α-II -spectrin, (a non-erythrocytic neuronal cytoskeletal protein with prominent axonal expression), and Glial Fibrillary Acid Protein, ([GFAP], an intermediate filament protein with predominant astrocytic expression in the CNS), each associated with a neurodevelopmental phenotype (MIM-300802, MIM-613477, and MIM-203450 respectively) (Amberger, Bocchini, Schiettecatte, Scott, & Hamosh, 2015; Amberger, Bocchini, Scott, & Hamosh, 2019) were examined in our samples.

## Methods

Exosomal analysis was conducted in plasma obtained from schizophrenia patients and matched controls participating in studies conducted by the Schizophrenia Neuropharmacology Research Group at Yale (SNRGY). Study protocols adhered to the Declaration of Helsinki and were approved by the Yale University Human Investigations Committee.

### Participant Evaluations

All participants underwent detailed medical and psychiatric evaluations, and a structured diagnostic interview (SCID-IV) as part of the standard evaluation after written informed consent was obtained. Subject demographics, information on duration of illness and antipsychotic medication (in patients) and substance use histories were systematically collected. Psychotic symptoms were assessed using the Positive and Negative Syndrome Scale (PANSS). Recent use of tobacco/nicotine and cannabis was assessed using the timeline follow back scale (Robinson, Sobell, Sobell, & Leo, 2014). Current antipsychotic medication dose was calculated in Olanzapine equivalents using the methods proposed in Leucht et al (S. Leucht, Samara, Heres, & Davis, 2016).

### Blood processing details

Venous blood was drawn into EDTA-containing tubes and centrifuged at ∼3000 rpm for 10-15 minutes. Resulting plasma was aliquoted (∼0.5ml) and frozen to -80°C. De-identified plasma samples underwent exosomal analysis to the laboratory of Dr. T Mohanakumar under an MTA between Yale University and Dignity Health/St. Joseph’s Hospital and Medical Center, Phoenix, AZ.

### EVs isolation and characterization

Exosomes were isolated from plasma using a standard exosome isolation kit (4478360, Invitrogen; Thermo Fisher Scientific, Waltham, MA) followed by 0.2 micron filtration and size determination using NanoSight using standardized procedures previously described (Gunasekaran et al., 2017). The exosome pellet of size from 40-200 nm was dissolved in PBS, the concentration of the exosomes was determined using the NanoSight and protein was measured using BCA kit. Exosomes were lysed with RIPA buffer; protein concentration was measured prior to immunoblots. Exosomal protein concentration was measured by BCA kit and an equal amount of protein (20 μg) was loaded per well.

### Nanoparticle tracking analysis ([NTA] NanoSight)

Vesicles were analyzed by nanoparticle tracking using the NanoSight NS300 system (Malvern, Great Malvern, UK). Samples were administered and recorded under controlled flow, using the NanoSight syringe pump and script control system. For each sample, 5 videos of 60 seconds duration were recorded, with a 10-second delay between recordings, generating 5 replicate histograms that were averaged. Therefore, the typical number of completed tracks per sample was approximately 1200. The area under the curve was calculated using Prism-4 software version 4.03 (Graph Pad, San Diego, CA), to give average particle counts from these replicates. Particles ranging in size from 40-200 nm were considered exosomes consistent with previously published methods (Yu et al., 2018) (see sample image shown in Figure 1).

**Figure 1.**
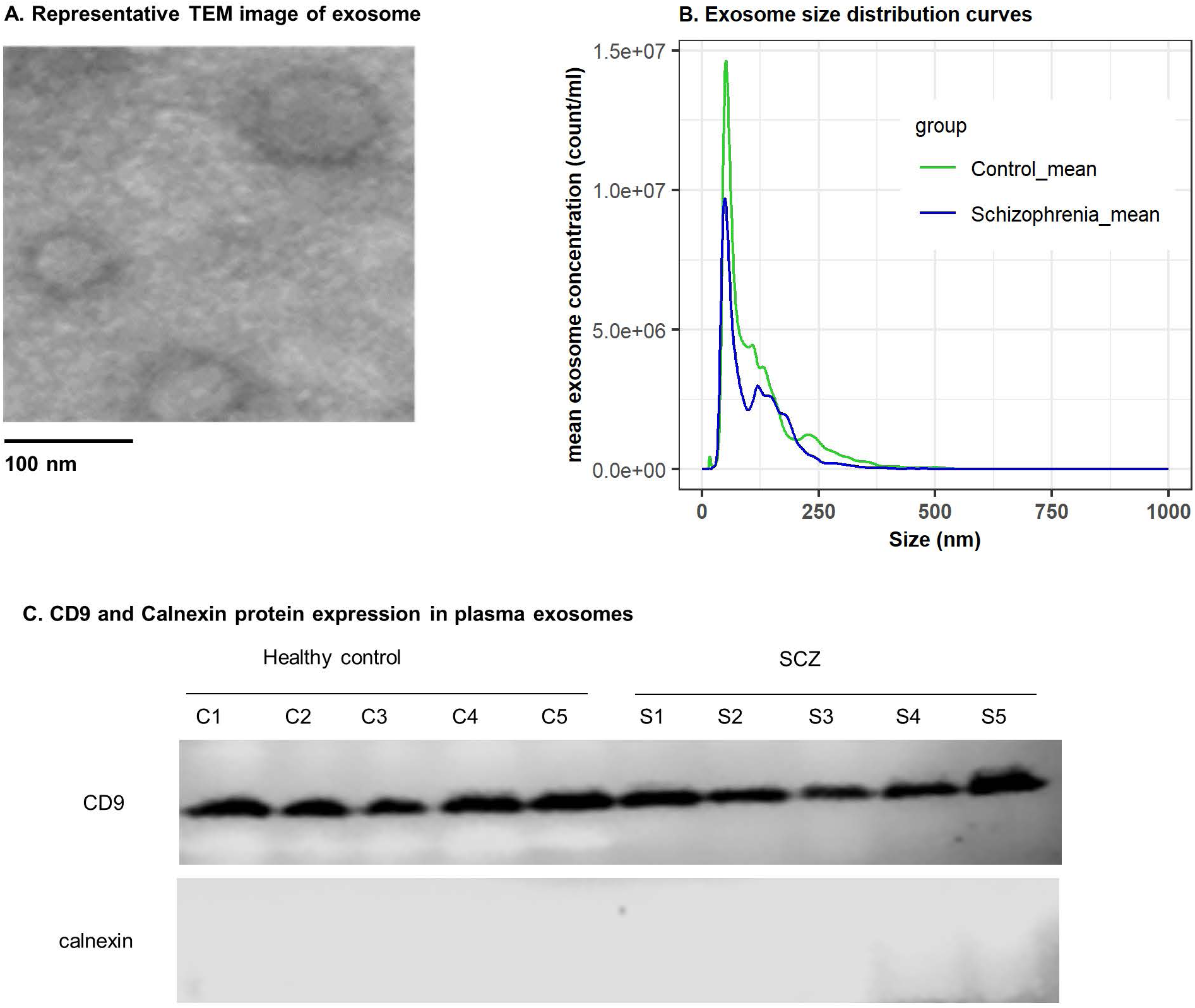
Characterization of plasma derived circulating exosomes with Transmission Electron Microscopy and Western blotting. a) Transmission Electron Microscopy image of a representative sample demonstrating that the isolates contained vesicles in the expected size range. b) The size distribution curves of mean exosome concentrations in schizophrenia and control groups. Both schizophrenia and control samples had maximal distribution and modal sizes of the isolated particles between 30-200 nm. The total concentration was lower in schizophrenia compared to controls. c) Western blot analysis with anti-CD 9 antibody in 5 representative cases and controls demonstrates that this constitutive exosomal membrane protein could be detected in all samples of schizophrenia and matched controls. Calnexin, a negative control protein for exosome secretory pathway was absent in all schizophrenia and control samples suggesting purity of exosome isolates.

### Western blot analysis

Western blotting was performed on the total protein extracts from exosome samples. RIPA lyses buffer (150 mM NaCl, 50 mM Tris, pH 8.0, 5 mM EDTA, 0.5% sodium deoxycholate, 0.1% SDS, 1.0% Nonidet P-40) with protease and phosphatase inhibitor cocktails (Sigma-Aldrich, St. Louis, MO, USA) was used for the total protein fraction. Protein concentrations in exosome extracts were determined using the BCA protein assay (Thermo Fisher). 20 μg of total lysates were diluted 1:1 in RIPA SDS-PAGE sample buffer, loaded onto polyacrylamide gels, and blotted onto polyvinylidene difluoride membranes (Bio-Rad). Membranes were blocked with 5% non-fat milk in PBS, pH 7.6, 0.2% Tween-20 for 1 hour and then incubated with mouse monoclonal anti-CD9 (1:1000, BD Pharmingan), mouse polyclonal anti-Synaptophysin (1:1000, Invitrogen), rabbit polyclonal anti-GFAP (1:1000, Invitrogen), rabbit polyclonal anti-α-II -spectrin (1:1000, Invitrogen), that binds α-II-spectrin and its break down products, and rabbit polyclonal anti-Calnexin (1:1000, Abcom) primary antibodies overnight. After washing in PBS-T, pH 7.6, 0.2% Tween-20, the membranes were incubated with a horseradish peroxidase-conjugated goat anti-rabbit antibody (1:2000, Cell Signaling) or anti-mouse antibody (1:2000, Cell signaling), and the immunoblots were visualized using ECL detection kits, (Pierce; Rockford, IL, USA). The band intensity was quantified using ImageJ software (National Institutes of Health, Bethesda, MD). CD9 is a tetra-spanin exosomal membrane protein marker that is typically absent in exosomes derived from B cells, NK cells and mesenchymal stromal cells and was used to confirm the identity of exosomes in the sample analyzed (Reyes, Cardeñes, Machado-Pineda, & Cabañas, 2018; Thery et al., 2018). Calnexin, a chaperone protein predominantly expressed in the endoplasmic reticulum, was included to demonstrate purity of the isolate from intracellular contaminants, specifically from the secretory pathway (Thery et al., 2018; Williams, 2006).

### Statistical Analysis

Demographic variables (age, gender and ethnicity) and substance use measures (nicotine and cannabis use) were summarized and compared for differences between the schizophrenia and the control groups using t-test and fisher’s exact text. In the schizophrenia group, PANSS measures and current antipsychotic exposure were summarized descriptively. The distributions of exosome concentrations in the two groups were examined with boxplots and compared with Mann-Whitney U (Wilcoxon rank sum) test. Exosome size distribution was plotted and examined using a density plot. The quantity of the three target proteins determined by densitometry were normalized to the quantity of CD9 in each sample. Due to non-normal distribution of protein quantities, group differences were examined with non-parametric Mann-Whitney U (Wilcoxon rank sum) test. Bonferroni correction was applied to adjust the p values for the independent comparisons of the three protein markers. The association of clinical and demographic variables with the concentrations of exosome target proteins were examined with Spearman’s correlation for continuous variables and Man-Whitney U test for binary variables. For age and sample age, partial correlations were calculated accounting for the effect of group. The association of lifetime cannabis use with protein concentration was examined separately in the case and control groups due to differing rates of use among schizophrenia and controls.

Variables with a significant association with protein concentrations were further examined in a generalized linear model to examine the main effect ‘group’ (schizophrenia vs controls) after adjusting for potential confounds. Data analysis was carried out in R version 3.6.1 (R Development Core Team, 2019) and the figures were generated with the packages tidyverse (Hadley Wickham, 2019) and ggpubr (Kassambara, 2019).

## Results

### Sample characteristics (Table 1)

The sample consisted of 24 (20 males;4 females) individuals with schizophrenia and 12 (8 males; 4 females) healthy controls. Demographic details and illness variables are presented in Table 1. The schizophrenia and the control groups were comparable in age, gender and ethnicity. A significantly higher proportion of participants with schizophrenia compared to controls had recent (past 30 days) cannabis. Three healthy controls had lifetime, but no recent history of exposure to cannabis, and none were current users of tobacco. None of the control participants had any major psychiatric or medical illness. Schizophrenia patients were ill for a median (IQR) duration of 2.65 (5) years and ∼80% of these participants were within the first six years of their illness. They were moderately to severely ill with a mean PANSS total score of 79.1 (Stefan Leucht et al., 2005) (Table 1). All except two patients were on oral or depot antipsychotic medications at the time of sample collection with a mean (SD) dose of 10.73 (9.66) mg in olanzapine equivalents (S. Leucht et al., 2016).

**Table 1:** Demographic and clinical details of the sample.

|  | Schizophrenia<br>(n = 24) | Control (n = 12) | Statistic | P value* |
| --- | --- | --- | --- | --- |
| Age in years – mean (SD) | 28.29 (9.23) | 27.16 (4.76) | t = 0.48 | 0.63 |
| Gender – F:M | 4:20 | 4:8 | OR (CI) =<br>2.43 (0.36 –<br>16.79) | 0.39 |
| Ancestry – n (%) |  |  |  | 0.58 |
| Caucasian | 7 (29.17%) | 5 (41.67%) |  |  |
| African American | 11 (45.83%) | 6 (50.0%) |  |  |
| Other | 6 (25.0%) | 1 (8.33%) |  |  |
| Current nicotine use– n (%) | 13 (54.2%) | 0 (0%) | OR > 2.44 | 0.002 |
| Recent cannabis use (Past 30 days)– n (%) | 7 (29.2%) | 0 (0%) | OR > 0.81 | 0.07 |
| Lifetime cannabis use – n (%) | 16 (66.7%) | 3 (25%) | OR (CI) =<br>5.68 (1.04 –<br>41.95) | 0.03 |
| Duration of illness in years – median (IQR) | 2.65 (5) | NA |  |  |
| PANSS positive - mean (SD) | 20 (3.43) | NA |  |  |
| PANSS negative - mean (SD) | 19.7 (4.12) | NA |  |  |
| PANSS general - mean (SD) | 39.4 (5.87) | NA |  |  |
| PANSS total - mean (SD) | 79.1 (9.03) | NA |  |  |
| Current anti-psychotic exposure in Olanzapine equivalents in mg/day mean (SD) | 10.73 (9.68) | NA |  |  |
| Current antipsychotic medication – n |  | NA |  |  |
| Risperidone | 6 | NA |  |  |
| Olanzapine | 4 |  |  |  |
| Aripiprazole | 4 |  |  |  |
| Quetiapine | 4 |  |  |  |
| Fluphenazine | 3 |  |  |  |
| Perphenazine | 2 |  |  |  |
| Paliperidone | 1 |  |  |  |
| Clozapine | 1 |  |  |  |
| Haloperidol | 1 |  |  |  |
| None | 2 |  |  |  |

\*Difference in age between the groups was examined with t-test. Gender, ancestry, smoking and cannabis use were compared with Fisher's exact test of independence.

### Exosome morphology, concentration, and size distribution (Figure 1)

Transmission electron microscopy (TEM) confirmed the morphology of the exosomes as spherical membrane bound vesicles with an approximate diameter of 100 nm (Figure 1a). All 36 samples of exosome isolates stained positively on immunoblotting with anti-CD9 antibody. None of the samples were positive with anti-calnexin antibody, demonstrating sample purity (Figure 1c) (Thery et al., 2018). The median (IQR) concentration of CD9 was 3272 (3657.79) in schizophrenia and 2300.52 (1974.34) in controls with no statistically significant difference between the two groups (Mann-Whitney U = 119, p = 0.42) (Figure 3a). There was also no statistically significant difference in the median concentration (case vs control, median (IQR) 6.49 *10^8^ (7.12*10^8^) vs 8.02 *10^8^ (3.4*10^8^)/ml, p = 0.07) or the modal size (case vs control, median (IQR) 85 (118) vs 51.5 (4.25) nanometers, p = 0.08) of plasma exosomes between cases and controls. The concentration and size distribution curves for schizophrenia and control subjects (Figure 1b) were within the ranges reported in the literature for circulating exosomes (Soares Martins, Catita, Martins Rosa, O, & Henriques, 2018).

**Figure 2.**
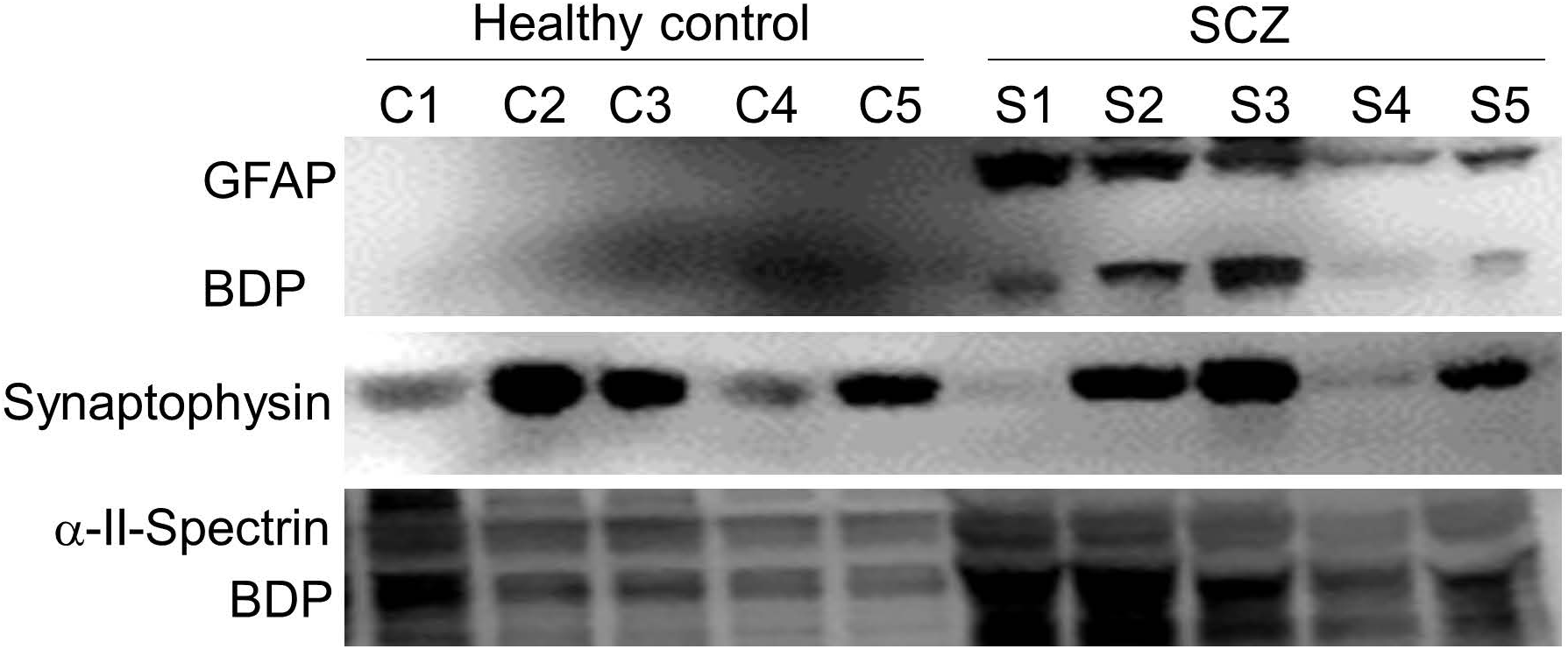
Western blot images of five representative samples from the schizophrenia and the control groups for three target proteins examined. Glial Fibrillary Acidic Protein (GFAP) and its break down products (BDP) was present in the exosomes derived from every sample in the schizophrenia group and was weakly positive only in 2 of the 12 control subjects tested (weak positive result for C4 shown in the image). Synaptophysin and α-II Spectrin with its BDP were detectable in samples from both schizophrenia and control groups.

**Figure 3.**
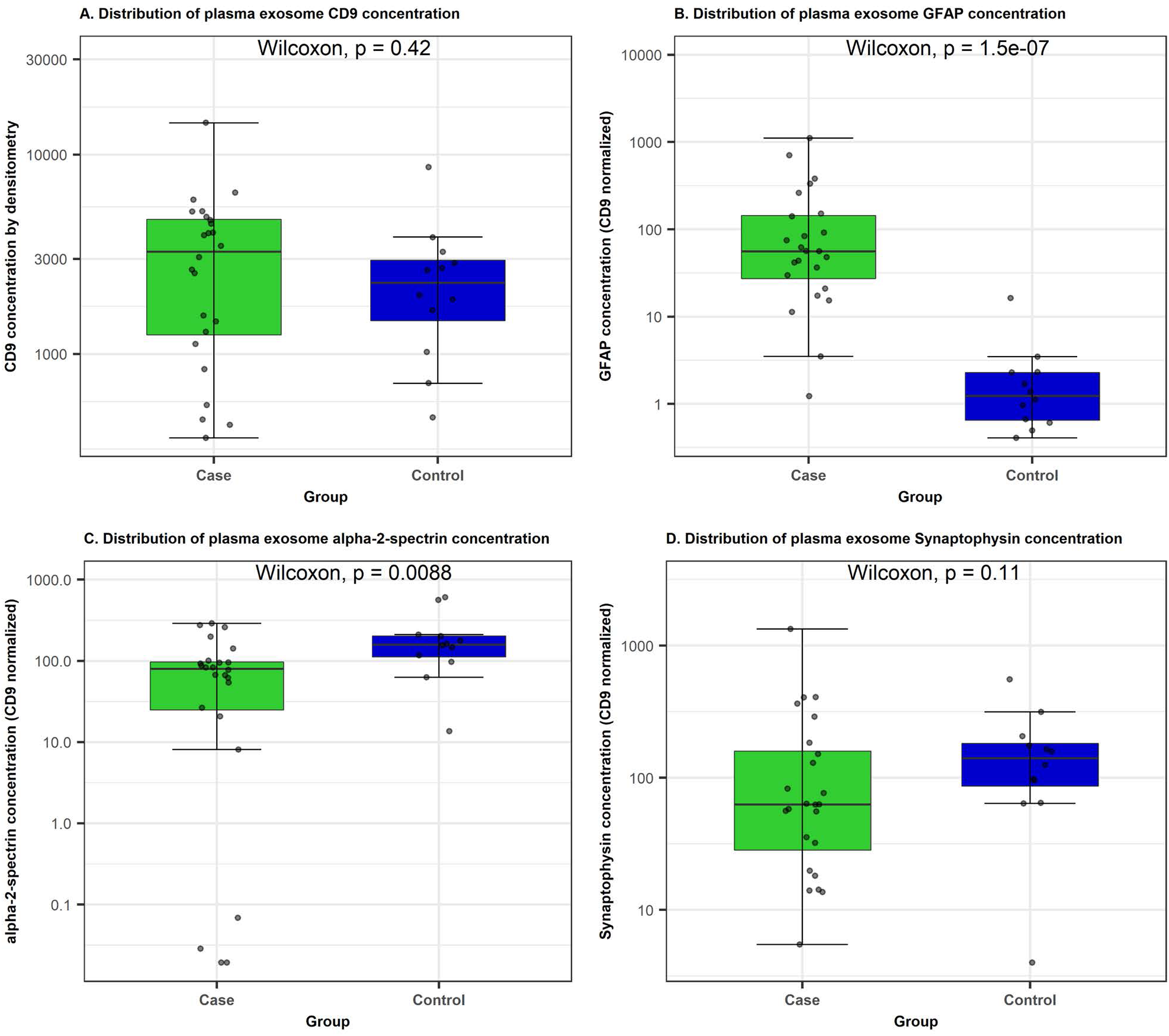
Box plots representing the differences in the concentration of CD9 and the three target proteins normalized to CD9 concentration in schizophrenia and control samples. Non-parametric Wilcoxon rank sum (Mann Whitney U) tests were used to compare the distributions in cases and controls due to non-normal distribution and extreme values. There was no significant difference in the concentration of CD9 between groups. GFAP was significantly elevated in schizophrenia and control samples. α-II Spectrin was decreased in cases compared to controls. Four of the 24 schizophrenia samples were very weakly positive for this protein on the western blots and are seen as outliers in the analysis. There was no statistically significant difference in the concentrations of synaptophysin. Y axis is presented in logscale.

### Astrocytic and neuronal cargo proteins in peripheral exosomes and relationship to clinical variables (Figure 2 and 3 and Table 2)

Figure 2 illustrates western blot images for a subset of schizophrenia and control samples for the three cargo proteins tested, GFAP, α-II-spectrin and synaptophysin. In qualitative analyses all 24 (100%) samples from schizophrenia subjects demonstrated positive immunostaining with polyclonal anti-GFAP antibody. In contrast, only 2 (16.67%) of the control samples showed GFAP immunopositivity. The CD9 normalized concentrations of GFAP was significantly higher in schizophrenia compared to healthy controls (median (IQR) 56.07 (126.02) vs 1.24 (1.68), Mann-Whitney U = 9, p = 1.5*10^−7^, p_corr_ = 4.5*10^−7^) (Figure 3b).

**Table 2:** Relationship between demographic and clinical variables and plasma exosome protein concentrations.

|  |  | **GFAP** | **Alpha-2-Spectrin** | **Synaptophysin** |
| --- | --- | --- | --- | --- |
| Age - r (p value) |  | -0.12 (0.53) | -0.23 (0.12) | -0.11 (0.54) |
| Gender – U (p value) |  | 79 (0.21) | 177 (0.01) # | 114 (.08) |
| Sample age – r (p value) |  | 0.047 (0.79) | 0.323 (0.06) | 0.054 (0.76) |
| Lifetime cannabis use - U (p value) | Schizophrenia | 79 (0.38) | 60 (0.83) | 61 (0.88) |
|  | Control | 8 (0.37) | 15 (0.86) | 18 (0.48) |
| **Variables examined only in schizophrenia sample** |  |  |  |  |
| Duration of illness – rho (p value) |  | 0.009 (0.97) | -0.22 (0.31) | -0.01 (0.96) |
| Current nicotine use – U (p value) |  | 38 (0.06) | 24 (0.006) # | 28 (0.012) # |
| Recent cannabis use – U (p value) |  | 64 (0.8) | 61 (0.95) | 54 (0.76) |
| PANSS positive – rho (p value) |  | -0.08 (0.7) | .27 (0.12) | -0.03 (0.87) |
| PANSS negative – rho (p value) |  | -0.04 (.84) | -0.22 (0.31) | -0.15 (0.48) |
| PANSS general – rho (p value) |  | -0.18 (0.4) | 0.5 (0.01) # | -0.23 (0.17) |
| PANSS total – rho (p value) |  | -0.08 (0.71) | 0.35 (0.09) | -0.16 (0.46) |
| Antipsychotic exposure in Olanzapine equivalents |  | -0.39 (0.06) | -0.39 (0.06) | -0.29 (0.17) |

Age, gender, cannabis use, and the duration of the sample storage did not have a significant association with GFAP concentration in the whole sample. Within the schizophrenia group, current psychopathology (PANSS total and subscale scores) did not have a significant association with GFAP concentration while antipsychotic dose had a negative but non-significant association suggesting that greater exposure to antipsychotics may be associated with a *lower* concentration of GFAP in circulating exosomes. Current nicotine use was present in 54.2% of the schizophrenia patients and none of the controls. Within the schizophrenia sample, nicotine use had a negative, but trend level association with GFAP concentration, such that nicotine users [median (IQR)] 43.6 (50.39) had a *lower* GFAP concentration compared to non-users 151.68 (352.2), Mann-Whitney U = 38, p = 0.06) α-II Spectrin immunopositivity was detected in all samples from schizophrenia (100%) and controls (100%) in qualitative assays. CD9 normalized α-II Spectrin concentration was significantly lower in schizophrenia samples, [median (IQR) 79.98 (77.27)] compared to controls [median (IQR) 158.8 (105.91)] and the difference was statistically significant (Mann-Whiney U = 221, p = 0.009, p_corr_ = 0.027) (Figure 3c).

Gender (Mann Whitney U = 177, p = 0.01), but not age (r_part_ = -0.23, p = 0.12), was associated with α-II Spectrin concentration, such that higher levels were observed in women (Table 2). Of the illness variables, PANSS general scores had a positive and antipsychotic dose had a negative but non-significant association with α-II Spectrin suggesting that greater exposure to antipsychotics may be associated with a *lower* α-II Spectrin concentration. Current nicotine users (within the schizophrenia group) had a significantly *lower* α-II Spectrin concentration [median (IQR) 53.81 (75.91)] compared to non-users [median (IQR) 95.91(175.67)] (Mann-Whitney U = 24, p = 0.006). The effect of gender and nicotine on α-II Spectrin concentration differences was examined with generalized linear model. The main effect of group continued to remain statistically significant (Beta = -1.27, SE = 0.43, p = 0.003) after adjustment for gender (Beta = - 0.056, p = 0.84) and current nicotine use (Beta = 0.82, p = 0.01) in the model, demonstrating lower α-II Spectrin concentration in schizophrenia compared to controls. Within the schizophrenia group, four patient samples had very low α-II Spectrin concentrations (Figure 3c). Group differences remained significant even after excluding these four samples and no differences were observed in the demographic or clinical variables in these four patients.

Synaptophysin immunopositivity was noted in all 24 samples from schizophrenia (100%) and in 10 out of 12 (83.3%) samples from control participants in qualitative assay. There was no statistically significant difference in the CD9 normalized concentration of synaptophysin between schizophrenia [median (IQR) 62.81 (153.45)] and control [median (IQR) 141.9 (125.93)] (Mann-Whitney U = 192, p = 0.11, p_corr_ = 0.32) (Figure 3d).

There was no significant association between demographic or clinical variables and synaptophysin concentration in the whole sample. Current nicotine users (within the schizophrenia group) had a lower concentration of synaptophysin [median (IQR) 55.55 (46.09)] compared to non-users [median (IQR) 184.42 (343.73)] that was statistically significant (Mann-Whitney U = 28, p = 0.012).

## Discussion

Our results demonstrate for the first time, a differential pattern of circulating exosomal proteins in schizophrenia compared to matched healthy controls. The ability to extract exosomes from plasma and identify neuropathology relevant signatures represents a significant advance in the quest for peripheral biomarkers of schizophrenia.

Specifically, we demonstrate higher concentration of GFAP, a well-recognized marker of astrocytic pathology (Kim, Healey, Sepulveda-Orengo, & Reissner, 2018), in plasma exosomes of schizophrenia participants in contrast to matched healthy controls. This finding is particularly significant given the absence of group differences in exosomal concentration and exosomal markers assessed in this study. All samples tested positive for CD9, an exosomal membrane protein, and negative for Calnexin, an endoplasmic reticulum protein demonstrating purity of the samples. Furthermore, exosomal samples from both groups were similar in concentration for synaptophysin, suggestive of the presence of neuronal derived exosomes (NDE) irrespective of the disease status. Thus, the significantly higher concentration of exosomal GFAP in schizophrenia samples is suggestive of *selective* enrichment of exosomal protein of astrocytic origin only in the patient samples.

These findings are consistent with the neuroinflammatory hypothesis and astro-glial pathology in schizophrenia (Dietz, Goldman, & Nedergaard, 2019; Kim et al., 2018). GFAP is highly expressed in astrocytes in the CNS, in a cell type specific fashion with minimal expression noted outside the CNS (Yang & Wang, 2015). Elevated GFAP expression is associated with increased astrocytic activation and inflammation (Kim et al., 2018). Post mortem studies in schizophrenia demonstrate that GFAP is upregulated in the pre-frontal cortex in schizophrenia although results from other brain regions remain mixed (Barley, Dracheva, & Byne, 2009; B. Farnsworth et al., 2017; Feresten, Barakauskas, Ypsilanti, Barr, & Beasley, 2013; Steffek, McCullumsmith, Haroutunian, & Meador-Woodruff, 2008; Toro, Hallak, Dunham, & Deakin, 2006). These mixed results may be related to region specific differences in GFAP expression, diversity of astrocytes with respect to GFAP positivity and heterogeneity within schizophrenia (Kim et al., 2018). For instance, Catts et al reported that elevated levels of brain GFAP and astrogliosis was observed only in a subset of schizophrenia patients and associated with greater neuroinflammation (Catts, Wong, Fillman, Fung, & Shannon Weickert, 2014).

Further studies are needed to determine if distinct patterns of exosomal GFAP expression are detectable in subsets of patients with schizophrenia. To our knowledge, there has been only one published study of free (non-exosomal) GFAP in CSF and serum samples schizophrenia, with no differences reported between first episode psychosis patients and controls (Steiner, Bielau, Bernstein, Bogerts, & Wunderlich, 2006). Exosomal analyses may offer greater ability to identify neuropathology relevant proteins given the cell-of-origin specificity of exosomes, and the enrichment and stability of their cargo protein.

GFAP expression may be affected by age (Catts et al., 2014; Rosengren, Wikkelso, & Hagberg, 1994) and antipsychotic medications (D. L. Farnsworth, 1976) such that brain tissue, CSF and serum GFAP increases with age (Rosengren et al., 1994), with very low levels in healthy individuals till around 40 years of age. The absence of detectable GFAP amongst our control subjects (mean age 27.16 years) is consistent with this literature. In contrast, the uniform GFAP positivity in our *age-matched* schizophrenia samples is particularly noteworthy. Amongst patients, there was no association of GFAP concentration with duration of illness, illness severity, substance use or exposure to antipsychotics. The existing literature on the impact of antipsychotic medications on GFAP expression is mixed (D. L. Farnsworth, 1976) and all patients had prior exposure to antipsychotics although two were currently not exposed. The non-significant trend for antipsychotic exposure to be associated with lower GFAP levels is noteworthy, given that the direction of association cannot explain the higher GFAP levels amongst patient samples. Current nicotine use was present only amongst patients with schizophrenia in this study in whom, current use was associated with lower GFAP levels. Similar to antipsychotic exposure, the directionality of the association is opposite to the observed group differences between schizophrenia and healthy controls and thus, cannot explain the observed group differences.

α-II Spectrin concentration was lower in schizophrenia compared to controls. α-II Spectrin levels were higher in women and within the schizophrenia group lower with antipsychotic exposure and current nicotine use. α-II Spectrin is a cytoskeletal protein expressed in non-erythroid tissue and enriched in the brain where it is present abundantly in neurons, particularly in axons and synaptic cytoskeleton (Pineda et al., 2007). α-II Spectrin and its break down products are detected in the cerebrospinal fluid as freely soluble proteins as well as in exosomes in patients with TBI and sub-arachnoid hemorrhage(Lewis et al., 2007; Manek et al., 2018; Pineda et al., 2007) suggestive of ongoing neuronal loss. Lower α-II Spectrin concentration in schizophrenia may be consistent with neuronal loss in this illness.

Synaptophysin expression was observed in all exosomal samples in our study and quantitative analyses revealed no group differences in levels. Similar to α-II Spectrin, synaptophysin levels were lower in current nicotine users. Synaptophysin is a synaptic vesicle membrane protein expressed in neurons and neuroendocrine cells where it plays a critical role in the formation and regulation of synaptic vesicles (Kwon & Chapman, 2011). Several studies have demonstrated reduced synaptophysin mRNA and protein expression in post mortem tissue of schizophrenia patients consistent with reduced synaptic vesicle density and function in schizophrenia as previously reviewed (Egbujo, Sinclair, & Hahn, 2016; Osimo, Beck, Reis Marques, & Howes, 2019).

Exosomal synaptophysin has also been examined in several neurodegenerative disorders such as Alzheimer’s disease(Goetzl, Kapogiannis, et al., 2016) and TBI(Manek et al., 2018). Goetzl et al reported lower levels of synaptophysin in NDE in frontotemporal dementia and Alzheimer’s disease and the reductions in synaptophysin significantly predated the cognitive decline(Goetzl, Kapogiannis, et al., 2016). Our study did not isolate NDE in plasma samples, although synaptophysin positivity in our samples is indicative of NDE content in our analytes.

### Influence of clinical variables on exosomal protein levels

In a cross-sectional study such as this, demographic/clinical variables must be considered. In our sample, the two groups were matched for age, gender, sample processing and duration of sample storage. There was no association between age and exosomal protein concentrations, but in this small sample, female gender was associated with higher levels of α-II Spectrin. The two groups were not matched for illness variables including psychotic symptoms, antipsychotic medications and current nicotine use, all of which were present in the schizophrenia group and not amongst controls. Other than a positive association between PANSS general score and α-II Spectrin concentration, there were no associations between symptoms of schizophrenia, duration of illness or exposure to antipsychotic medications and exosomal proteins assessed. Current nicotine use was associated with lower levels of all 3 exosomal proteins assessed, although the effect on GFAP was not statistically significant. Importantly the between group (schizophrenia vs. healthy control) differences in exosomal GFAP and α-II Spectrin levels remained significant even after controlling for nicotine use. The association between nicotine exposure and exosomal proteins warrants further study-especially given high rates of cigarette smoking in schizophrenia.

Our understanding of exosomal biology, cargo selection and function are still emerging and it is recognized that exosomes play critical roles in physiological and pathological processes in the CNS (Stahl & Raposo, 2019). Elevated exosomal GFAP in schizophrenia is consistent with greater neuroinflammation and astrocytosis-but may also represent reduced integrity of the blood brain barrier in the context of inflammation. However, a lack of group differences in exosomal numbers and synaptophysin concentration, and lower exosomal α-II Spectrin in schizophrenia, argues against the latter. α-II Spectrin, in addition to representing a neuronal/synaptic cytoskeletal protein is also an integral constituent of extracellular vesicles. Whether the reduced α-II Spectrin levels in schizophrenia reflects altered exosome production or secretion related to the illness remains unclear and needs further study. The pattern of elevated GFAP and reduced α-II Spectrin is consistent with increased astro-glial inflammation and neuronal loss in schizophrenia (Laskaris et al., 2016). Unlike exosomal GFAP, exosomal α-II Spectrin and synaptophysin were detected in both groups and may also be reflective of the expression of these two proteins in non-neuronal tissues, and future studies should specifically examine their expression in circulating NDE in schizophrenia.

### Strengths and Limitations

These data represent the very first study examining neuropathology relevant protein biomarkers in circulating plasma derived exosomes in schizophrenia. The analytical methods employed here were consistent with the recommended guidelines included in the “Minimal information for studies of extracellular vesicles 2018 (MISEV2018): a position statement of the International Society for Extracellular Vesicles and update of the MISEV2014 guidelines” for studies on exosomes (Thery et al., 2018). As recommended in the MISEV2018 guidelines, exosomes were extracted from plasma samples with a standardized protocol using exosome separation kits. The morphology of the exosomes was characterized with both TEM and NTA. NTA was also used to determine the concentration per ml of plasma sample. Exosomal confirmation included assessment of transmembrane CD9 protein by Western blot. Further, the purity of the isolate was demonstrated with the absence of calnexin protein in the sample, thus precluding contamination from other particles in the secretory pathway such as endoplasmic reticulum. Finally, participants were well characterized for demographic and illness variables.

We did not specifically extract NDE from plasma samples in this study. Further, the lack of a second patient group prevents our ability to account for the effect of chronic illness or stress. Finally, we did not study antipsychotic naïve patients in order to examine the effect of any antipsychotic medication exposure on exosomal protein expression. These limitations notwithstanding, these data strongly support the potential of exosomal proteins as peripheral biomarkers of neuropathology in schizophrenia.

### Future Directions

These initial results provide the impetus to examine a wider array of exosomal derived proteomic signatures in schizophrenia. Whether the observed differences are also present in other psychiatric disorders remains to be examined. Finally, larger and longitudinal studies are needed to determine whether these findings represent trait markers of the disorder or vary by stage of illness, antipsychotic treatment, or other clinical factors in schizophrenia.

## Acknowledgements and funding

This work was supported by unrestricted research funds (SNRGY), Department of Veteran’s Affairs (Mohini Ranganathan), Brain and Behavior Research Foundation (Suhas Ganesh) and St. Joseph’s Foundation (T. Mohanakumar).

